# Combination Treatment Optimization Using a Pan-Cancer Pathway Model

**DOI:** 10.1101/2020.07.05.184960

**Authors:** Robin Schmucker, Gabriele Farina, James Faeder, Fabian Fröhlich, Ali Sinan Saglam, Tuomas Sandholm

## Abstract

The design of efficient combination therapies is a difficult key challenge in the treatment of complex diseases such as cancers. The large heterogeneity of cancers and the large number of available drugs renders exhaustive *in vivo* or even *in vitro* investigation of possible treatments impractical. In recent years, sophisticated mechanistic, ordinary differential equation-based pathways models that can predict treatment responses at a *molecular* level have been developed. However, surprisingly little effort has been put into leveraging these models to find novel therapies. In this paper we use for the first time, to our knowledge, a large-scale state-of-the-art pan-cancer signaling pathway model to identify potentially novel combination therapies to treat individual cancer cell lines from various tissues (e.g., minimizing proliferation while keeping dosage low to avoid adverse side effects) and populations of cancer cell lines (e.g., minimizing the maximum or average proliferation across the cell lines while keeping dosage low). We also show how our method can be used to optimize the mixtures and dosages used in *sequential* treatment plans—that is, optimized sequences of potentially different drug combinations—providing additional benefits. In order to solve the treatment optimization problems, we combine the Covariance Matrix Adaptation Evolution Strategy (CMA-ES) algorithm with a significantly more scalable sampling scheme for truncated Gaussian distributions, based on a Hamiltonian Monte-Carlo method. These optimization techniques are independent of the signaling pathway model, and can thus be used for other signaling pathway models also, provided that a suitable predictive model is available.

## 1. Introduction

Rational design of combination therapies is a difficult but important challenge in the treatment of complex diseases such as cancers [36, 33, 2]. The large heterogeneity of cancers and number of available drugs renders exhaustive *in vivo* or even *in vitro* investigation of treatments impractical. Accordingly, computational models that enable—even individualized—prediction of drug sensitivity have to be employed [42]. To this end, sophisticated mechanistic, ordinary differential equation (ODE) models for sensitivity prediction have been developed [6, 13, 38, 53, 8, 29, 41]. However, so far little effort has been put towards using these models to actually design treatments. Typically, only the temporal aspect of when to administer drugs [45, 51] is considered, but not which drugs to pick.

In this paper we present a methodology for *in silico* combination treatment optimization which leverages a large-scale mechanistic pan-cancer pathway model [13]. A robust evolutionary optimization algorithm is modified and used to guide the search for effective drug combination. The proposed framework can be easily adapted to find treatments for other complex diseases than cancers, as long as a suitable predictive model is available. Our experiments show how the approach can lead to effective combination therapies—trading off low proliferation with adverse side effects—targeting single cancer cell lines or multiple-cell lines at once. Furthermore, we show how our method can be used to optimize *sequential treatment plans* which apply varying drug cocktails in sequence.

To our knowledge, this is the first application of a large-scale pan-cancer pathway model to discover novel combination therapies. We adapt non-convex optimization techniques and use an efficient parallelization scheme which enables experiments on dozens of cell lines and combinations of 7 anti-cancer agents at low cost. Three different treatment scenarios targeting single as well as multiple cell lines at once are formalized as optimization problems and experimental studies are conducted. Our simulations led to the discovery of novel treatment approaches in the form of drug combinations that achieve better predicted treatment effects at lower concentrations than the best prior approaches.

### Related Work

The use of mathematical modeling for the design of cancer therapies has a rich history. Early studies combined optimal control theory with a growth model of bone cancer to find treatment regimes which balance reductions in cell population with administered dosage of a single drug [3, 48]. Moreover, evolutionary game theory [46, 27] was used to analyze the adaption of cell populations under selective pressure, especially with regards to population size [23, 16, 10], and emergence of drug resistance [34, 49, 39]. Sandholm [44, 43] proposed modeling treatment planning— and steering biological entities more generally—as a multi-step game between a biological entity and a treater, for the purposes of computationally constructing steering plans that can involve combination therapies, sequential plans, and conditional plans (aka. adaptive treatments). He proposed modeling the biological entity in the game 1) using a behavioral model if there is enough data, 2) as a game-theoretic worst-case adversary if there is not enough data, or 3) as an opponent with limited lookahead so it can be exploited by luring it into traps. (Specific algorithms have since then been developed for exploiting an opponent’s limited lookahead in imperfect-information games [31, 30], but they have not yet been applied to biological settings.) In that taxonomy, the present paper falls under approach (1). Adaptive treatment regimes [15]—that is, regimes that monitor tumor development and use predictive models to adapt reactively—have led to promising preclinical trials on breast cancer [9] and Phase 2 clinical trials on prostate cancer [54]. Multiple *in vitro* studies [7, 14, 1] investigated the emergence of drug resistance and showed advantages of adaptive treatment regimes. A recent line of work [50, 52, 51] investigates benefits of combination treatments on the development of drug sensitivity. Stackelberg games [47] have also been proposed for treatment design.

While these prior approaches rely on rather high-level abstractions of the underlying biology, our work employs a detailed, mechanistic pan-cancer signaling pathway model [13]. It can be individualized to cell-lines using sequencing data, which is important to account for heterogeneity in response. It describes the action of 7 small molecule inhibitors, which enables the design of higher order combinations. Furthermore, previous evaluations of the model indicated that it is capable of quantitatively accurately predicting the effect of drug combinations from single drug treatments [13], which is essential for the reliability of treatment strategies we propose. The only prior work [32] in this direction uses a Boolean T-cell signaling pathway model [25] which yielded—due to its Boolean nature—mainly qualitative insights. Our work serves as a proof of concept of how biologically accurate quantitative signaling pathway models can be combined with optimization algorithms to discover effective combination therapies, including multi-step ones. Our methodology and computational approach enabled us to perform extensive experiments with combinations of 7 existing anti-cancer agents on dozens of cancer cell-lines yielding promising directions for future laboratory studies.

## 2. Methods

In this section we present our approach in detail. We first discuss the pan-cancer signaling pathway model that is used to predict treatment responses. Building on the predictions of this model three different combination treatment optimization problems are introduced. In order to tackle these problems we discuss modifications to the CMA-ES algorithm [22], to make it suitable for our domain. Finally, we discuss studied cell lines and combination treatments as well as implementation details.

### 2.1 Pan-Cancer Cell Simulation

For our treatment optimization experiments, we employed a pre-existing large-scale mechanistic pan-cancer signaling pathway model [13]. The model describes the effects of 7 targeted anti-cancer agents on multiple cancer-associated pathways at the molecular level as an ODE model. In total, the model describes the temporal development of 1228 different molecular species, that is, concentrations of ligands, protein complexes or drugs, through 2704 reactions using a total of 4104 parameters. Every model simulation reports a proliferation score

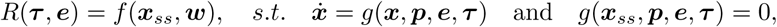

where 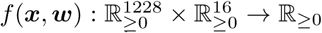 is a phenomenological function that maps molecular abundances to proliferation scores, ***x***_*ss*_ are molecular abundances defined by the steady state of the ODE model and 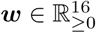 are mapping coefficients, which are free parameters of the mapping function. Here, 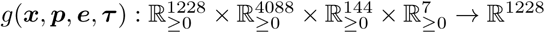 is the right hand side of the differential equation. 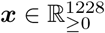, the kinetic parameters 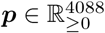 are biophysical rate constants such as binding rates or catalytic activities, which are free parameters for the differential equation model 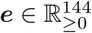 are mRNA expression levels for 108 different genes and 36 gain of function mutations described by the model, which can be used to individualize the model to specific cell lines 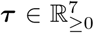are drug concentrations, which define the concentrations of individual drugs in the extracellular compartment.

To be biologically meaningful, the proliferation score *r* has to be normalized to the proliferation score for the untreated condition with ***τ*** = 0. The normalized relative proliferation score *V* (***τ***, ***e***) = *R*(***τ***, ***e***)*/R*(0, ***e***) can be directly compared to experimental observations from cell viability assays such as CellTiter-Glo [20], which quantify the difference in cell counts between treated and untreated conditions, thus accounting for the net sum between cell growth and cell death. For all simulations, we used previously reported values for ***p*** and ***w***, which were obtained by training the model on relative proliferation data from 120 cell lines from the Cancer Cell Line Encyclopedia [4]. We used the AMICI software package [12]—which internally uses CVODES [26]—to solve the differential equation model (that is, the signaling pathway network variables) to steady state after each treatment. Default AMICI integration and steady state tolerances were used.

### 2.2 Multi-Drug Treatment Optimization

We leverage the pathway model discussed in Section 2.1 to identify novel combination therapies for a variety of cancers using 7 preexisting drugs. Formally, we represent a multi-drug treatment by a 7-tuple 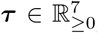 Entry *τ*_*i*_ is the concentration of the *i*-th drug contained in treatment ***τ*** In nanomoles (nM). Mathematically, the set of treatments considered in this paper is represented by 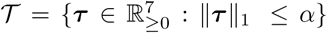 that is, the set of all combination therapies whose total dosage is below threshold value *α*. In prior work, the pathway model had been fitted with clinical data administering concentrations in the range from 2.5nM to 8000nM. Thus, we use a value of *α* = 8000 to ensure that the optimization domain 𝒯 resembles the training data in terms of total dosage.

An effective treatment needs to trade off between desired and adverse effects. For each cell line *c* the model defines a function *V*_*c*_:*𝒯 ⟶* R≥ *V* (***τ***, ***e*_*c*_**), which given a treatment *τ* ∈ 𝒯 and a vector of expression levels ***e*_*c*_**, predicts the relative proliferation value of *c* when subjected to *τ*. The predicted relative proliferation is used to capture desired treatment effects. Because the literature does not offer a concise way to quantify adverse effects on healthy cells caused by a combination of multiple drugs, we apply a mathematical regularization function *R* to the treatment vector as an idealized measure. Prior work has used L1 [3, 48] and L2 [35] regularization for this purpose. In our experiments we use L1, L2 and sum of logs regularization and compare differences in resulting treatments.

The following three subsubsections, respectively, introduce three different treatment optimization problem classes that are addressed in our simulation study.

#### 2.2.1 Optimizing the Single-Step Treatment of a Single Cell Line

First, we focus on identi-fying a treatment *τ* ∈ 𝒯 that is effective for a specific cell line *c*. An optimization problem which balances relative proliferation score and adverse effects is given by

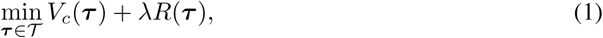

where the penalty parameter *λ* sets the weight of adverse effects as quantified by regularizer *R*. Large values of *λ* favor conservative treatments while low values favor more aggressive treatments.

#### 2.2.2 Optimizing the Single-Step Treatment of a Population of Cell Lines

Tumors often feature multiple sub-clones that feature different sets of mutations and expression levels. To avoid resistance, all sub-clones have to be targeted effectively. As a proxy for these sub-clones, we consider multiple cell lines with the same tissue of origin. Accordingly, we try to construct treatments *τ* ∈ 𝒯 that are simultaneously effective on a set of different cell lines *𝒞*. The optimization problem is

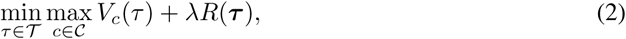

where the objective function only considers the highest predicted proliferation value following treatment *τ* among the cell lines 𝒞 in, that is, the most proliferated cell line. This objective favors treatments that reduce the proliferation values of all cell lines evenly.

An alternative is to use a weighted sum of the individual proliferation scores. This could be useful, for example, for finding personalized treatments when the distribution of cell types in a tumor is known. When starting weights are used, that objective function tries to minimizes the average proliferation of all cell lines in set *𝒞*. An experimental comparison of both approaches is provided in Appendix A.2.

#### 2.2.3 Optimizing Sequential Treatment Plans

For a heterogeneous population of cell lines, it may not always be possible to find a single treatment that is effective in all cell lines. To address this, we also investigate the discovery of a *sequential treatment plan*, that is, a sequence of combination treatments (***τ***_1_, …, ***τ***_*n*_) that is effective on a set of cell lines 𝒞. Let the space of sequential treatments 𝒯^*n*^ be the n-ary Cartesian power of the space of drug combinations 𝒯. A treatment plan optimization problem is now given by

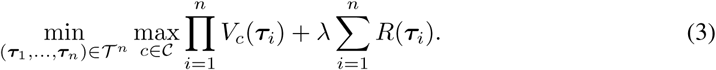

For each cell line 𝒸 ∈ 𝒞, the relative proliferation value is computed by taking the product of the predicted relative proliferation values at the individual treatment steps. This assumes that the growth of a cell line during one of the steps of the treatment plan multiplicatively affects the growth of that cell line in the next treatment step. A simple, biologically plausible model that satisfies this assumption is an exponential growth model with different, drug-dependent growth rates in each treatment step. Appendix A.1 discusses the exponential growth model in more detail.

Similar to the multi-cell line setting, this objective function considers the highest proliferation value to find a therapy that is effective for all 𝒸 ∈ 𝒞. The advantage of sequential plans compared to *time-invariant plans* is that the use of multiple specialized drug-combinations targeting different subsets 𝒞 of one at a time can be more effective than a single general *τ* ∈ 𝒯 targeting all of cell lines at once. A small illustrative example for this is shown in Figure 1. In this paper discrete 72h time steps are naturally enforced in that the path-way model is simulated from one steady state to the next.

**Figure 1:**
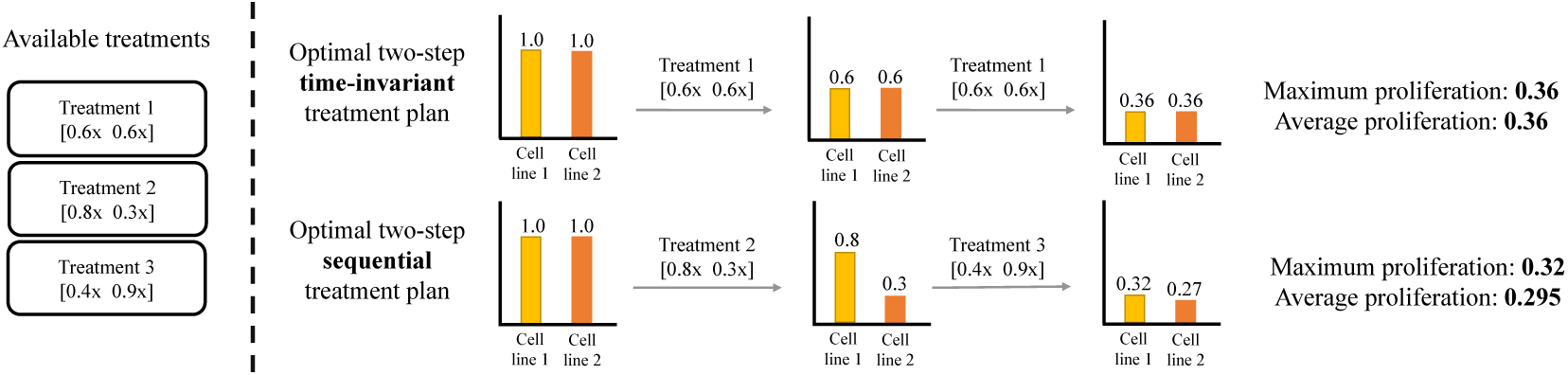
A potential benefit of a *sequential treatment plan* compared to a *time-invariant treatment plan* is that the use of different specialized drug-combinations (targeting fewer cell lines at once) allows for more effective therapies. In the illustrative example above with two cell lines and three available treatments, the optimal two-step time-invariant treatment leads to a relative proliferation score of 0.36 for both cell lines. Meanwhile, the optimal two-step sequential treatment plan achieves relative proliferation scores of 0.32 and 0.27 for cell lines 1 and 2, respectively.

### 2.3 Optimization Process

The deployed pathway model behaves in a non-convex way when interpolating between drug combinations. Because of this the optimization problems proposed in Section 2.2 are non-convex and there is no known algorithm that is both scalable and guaranteed to find an optimal solution in every case.

In this work we implemented *covariance matrix adaption evolution strategy* (CMA-ES)—a robust and sample-efficient algorithm [22]. The underlying idea of CMA-ES is to iteratively generate a set of solution candidates whose objective scores are then evaluated. After this, a number of *elites*—that is, the solution candidates with the best objective scores—are selected which are then used to generate the solution candidates for the next iteration step. The CMA-ES algorithm does this by maintaining a mean vector and covariance matrix describing a multivariate Gaussian distribution. At each step, solution candidates are sampled and elites are selected to update the mean and covariance matrix in a way that increases the likelihood of reaching previous elite solution candidates.

Over the years, a large variety of CMA-ES variations have been proposed and applied to various domains. Our implementation of the algorithm exactly follows that presented in [28]. However, we had to make certain modifications to that algorithm to account for the fact that the domain of treatments 𝒯 is a constrained set. We will discuss those modifications next.

### 2.3.1 Sampling from a Constrained Space

During the sampling step, CMA-ES generates a set of solution candidates by sampling from a multivariate Gaussian distribution. When dealing with a constrained domain, naive sampling can lead to the generation of infeasible solution candidates. A popular way to deal with this problem is to simply reject the infeasible points and to sample again until all candidates are feasible [21, 5]. This process effectively transforms the multivariate proposal distributed into a truncated Gaussian.

However, this approach fails in our treatment domain. 𝒯 The volume of domain roughly shrinks with a factor 1*/d*!, where *d* is the problem dimension. With increasing dimensionality, the vast majority of sampled solution candidates needs to be rejected, rendering the naive rejection-based approach infeasible. To avoid this issue, we employ a Hamiltonian Monte Carlo method [40], which can directly generate samples from a truncated multivariate Gaussian distribution that can be constrained by linear and quadratic inequalities. Even for lower-dimensional problems, this method speeds up the sample generation process by multiple orders of magnitude. Without this modification optimization of *n*-step sequential treatment plans (*d* = 7*n*) would not have been possible.

### 2.4 Cell Lines, Penalties, and Reference Drug Combination used in the Experiments

We experiment with 12 colorectal, 19 melanoma, 10 pancreatic, and 20 breast cancer cell lines on which the model discussed in Section 2.1 was trained in prior work. Cancers from these tissues have a high frequency of BRAF and RAS mutations, for which a large fraction of drugs in the model is thought to be effective. We varied the penalty parameter *λ* from 10^−7^ to 10^*−1*^with exponent steps of 0.25 (0.05 for the sequential experiments). For every problem configuration, the optimization algorithm is initialized with 3 different random seeds and run for 400 iterations. The search result with best objective function value is reported. We compare the optimized treatments to two baselines. The first baseline is the best single-drug treatment which is determined as follows. For each of the 7 drugs, treatments using concentrations in the range from 0 nM to 8000 nM are considered. Their objective and relative proliferation values are evaluated at 1 nM steps. For a given penalty parameter *λ*, the best single drug treatment is identified by its objective value. The second baseline are two-drug combinations that use a mixture of PLX-4720 (RAFi)+PD0325901 (MEKi). PLX-4720 and PD0325901 serve as a proxy for the clinical grade combination therapy of Vemurafenib (RAFi) and Cobimetinib (MEKi) for BRAF mutant melanoma [33]. Vemurafenib is the clinical analogue of PLX-4720 and PD0325901 and Cobimetinib are allosteric inhibitors that target similar pockets in MEK molecules. As it was difficult to find precise information on the clinical mixture ratios for these two drugs, we consider ratios from 0%-100% evaluated at 5% steps. As for the single drug baseline, treatments that use a total concentration in the range from 0 nM to 8000 nM are evaluated at 1 nM steps, and the two-drug treatment that achieves the best objective value is used as the second baseline.

#### Computation

All experiments were conducted using a compute cluster. Each individual experiment was run on a single 64-core server with AMD Opteron(TM) 6272 2.1 GHz processors and required less then 64 GB of RAM. Each prediction of proliferation for a given cell line and treatment (that is, one call to the function *V*_*c*_) took about 1 second. This dominated the run-time of the CMA-ES algorithm. We parallelized the evaluation of treatment candidates generated by the CMA-ES algorithm, and furthermore, for each solution candidate, parallelized the evaluation of that treatment on the different cell lines. In this way, we were able to run all the experiments in less than two weeks.

## 3 Results

In this section we show the effectiveness of the treatments discovered by the modified CMA-ES algorithm introduced in Section 2.3 for the three settings proposed in Section 2.2. For each setting, the findings are illustrated and the resulting treatments are compared to the two baselines.

### Optimizing the Single-Step Treatment of a Single Cell Line

For the first experiment, the objective function defined in Section 2.2.1 is used to find effective drug-combinations for individual cell lines. Figure 2 visualizes the optimization results for K029AX—a melanoma cancer with BRAF V600E mutation—for three different types of regularization. The treatments discovered by the algorithm achieve significantly lower relative proliferation values at lower total dosage than the two baseline treatments. For low penalties all regularizations lead to similar treatment compositions. For higher penalties L2 regularization leads to treatments that use more drugs at lower dosage and logarithmic regularization leads to treatments that use fewer drugs at higher dosage. Logarithmic regularization penalizes combinations treatments harshly and a low objective value does not always identify a strong treatment. Further results for A2058, MDAMB435S—two other melanoma cancer with BRAF V600E mutation—are provided in Appendix A.4. For A2058 the optimized combination treatments are significantly more efficient than the baselines. For MDAMB435S, the clinical-grade combination therapy that uses PLX-4720 and PD0325901 is already very effective and the discovered treatment only leads to slight improvements.

**Figure 2:**
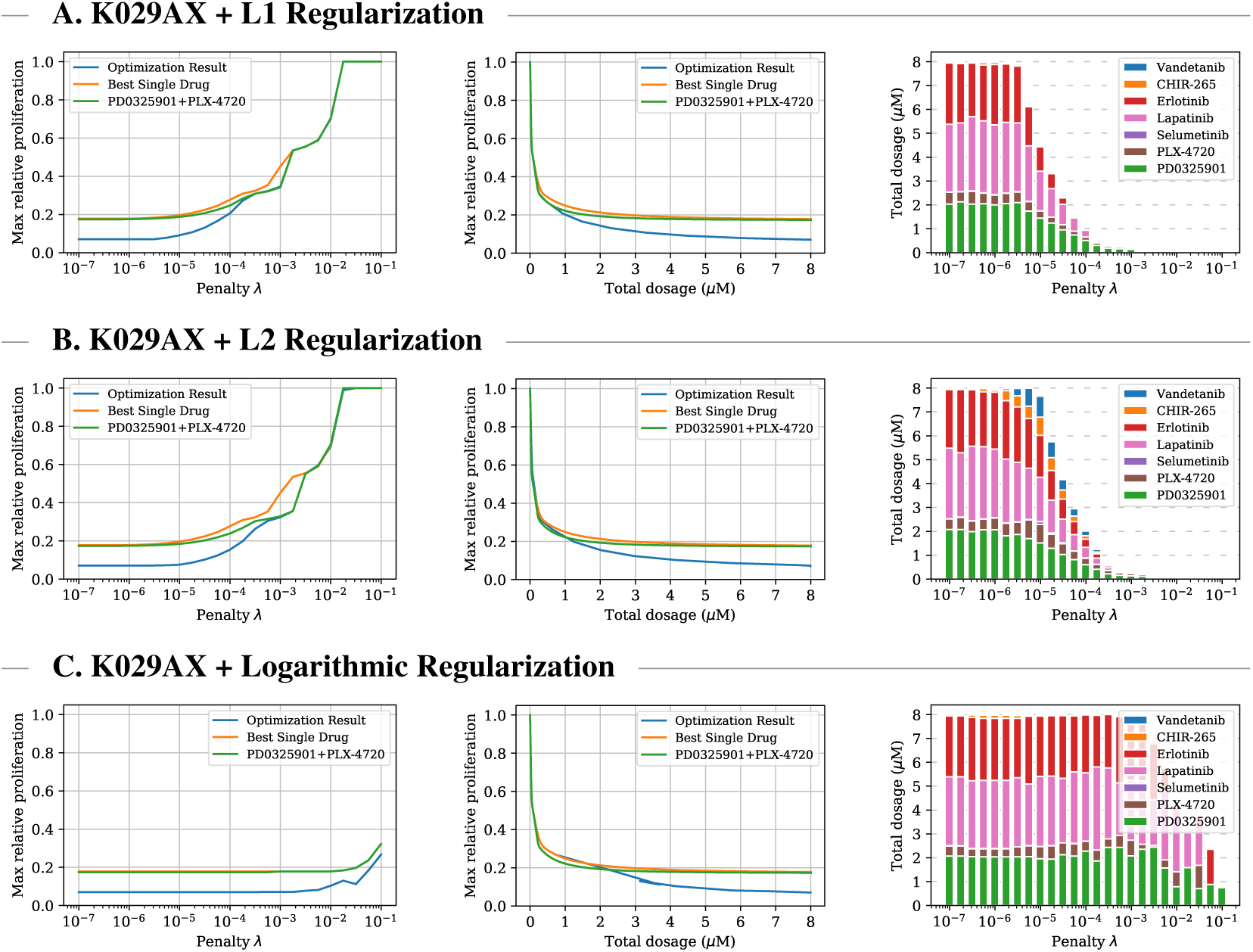
Comparison between optimized single-cell multi-drug treatment, optimal single-drug treatment, and optimal PD0325901/PLX-4720 combination treatment for K029AX—a melanoma cell line with BRAF V600E mutation—for three different types of regularization. Left plots: optimal treatment as identified by the objective function for different penalty parameters. The middle plots: relationship between administered total dosage and achieved proliferation value regardless of penalty and objective value. Right plots: composition of the multi-drug treatments. For all three types of regularization the optimization process leads to combination treatments which achieve significantly lower relative proliferation values at lower concentrations than single and two-drug treatment.

### Optimizing the Single-Step Treatment of a Population of Cell Lines

For the second experiment, the objective function defined in Section 2.2.2 is used to find drug combinations that minimize the maximum relative proliferation value predicted by the pathway model over sets of cell lines originating from skin (𝒞_Melanoma_), large-intestine (𝒞_Colorectal_), pancreas (𝒞_Pancreatic_), and breast (𝒞_Breast_) tissues, respectively. Findings for colorectal cell lines under L1 regularization are visualized in Figure 3. Experimental results for all tissues under all three types of regularization are provided in Appendix A.5. For all four tissues, the discovered treatments achieve significantly lower maximum relative proliferation values than the single-drug and PD0325901/PLX-4720 combination baselines at medium and high dosages. Especially for pancreatic cell lines, the optimized treatments reduce the cancer cell viability by a factor of up to three. For breast cancers, the optimization process leads to drug combinations that achieve notable treatment effects even at low dosage. We observed some variance in the optimized treatments when using low penalty values. We performed additional experiments with warm-starts and a PCA analysis which indicate that this behaviour only occurs under low penalties and that solutions are unique for medium and high penalties. It also empathized the issue of local minima under logarithmic regularization. More details are provided in Appendix A.3.

**Figure 3:**
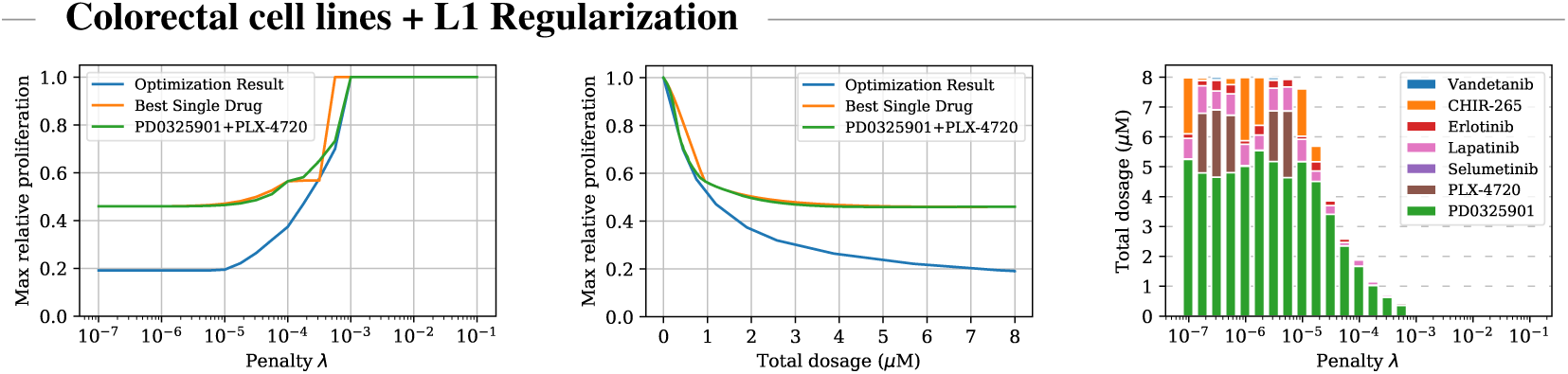
Comparison between optimized multi-cell combination, optimal single-drug, and optimal PD0325901/PLX-4720 combination treatment for colorectal cancers under L1 regularization. Left plot: optimal treatment as identified by the objective function for different penalty parameters. Middle plot: relationship between administered total dosage and achieved proliferation value regardless of objective values. Right plot: composition of the multi-drug treatments.

### Optimizing Sequential Treatment Plans

The third experiment investigates 2-step treatment plans and uses the objective function defined in Section 2.2.3 to find sequences of drug combinations that are effective on cell lines originating from the same tissue. In this setting we compare the performance of optimized sequential treatment plans (that is, ones that can use different drug combinations and dosages at the two treatment steps) against optimized time-invariant treatment plans (that is, ones that have to use the same drug combination and dosage in each of the two treatment steps). With these candidate drugs and cell lines, only very slight benefits were gained from allowing time-varying treatments. However, in a few cases at medium dosages we observed some larger gains. One example for an effective 2-step plan for colorectal cell lines under L1 regularization is shown in Figure 4 and Appendix A.6. A sequential plan that employs one aggressive drug-combination of PD0325901, PLX-4720, and Erlotinib and then a more conservative—that is, lower-dose—combination of PD0325901, Lapatinib, and Erlotinib achieves a maximum proliferation value of 0.6048 which is 13% lower than the proliferation value achieved by the optimized time-invariant treatment plan (0.6978), which uses a combination of PD0325901 and PLX-4720 at medium dosage.

**Figure 4:**
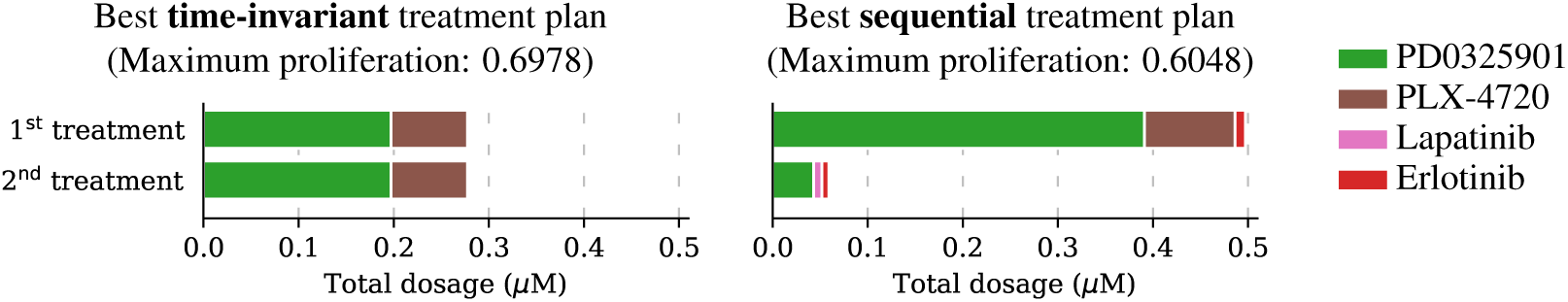
Comparison between drug cocktails administered by the optimized two-step treatment plan and the optimized two-step time-invariant treatment plan for colorectal cell lines under L1 regularization at the same total drug dosage (550 nM). The 2-step plan uses a high-dose treatment followed by a low-dose treatment. This achieves maximum proliferation 0.6048, which is more effective than the time-invariant treatment plan which only achieves 0.6978.

## 4 Discussion

Our approach discovered treatment strategies that deviate from current clinical first line treatment strategies. Without this result, the algorithm would not be of much use, as it would not predict anything new and just recapitulate what we already know. Yet, we have to carefully examine whether the proposed strategies are plausible from a biological perspective. For the investigations with BRAFV600E skin cancer cell lines, the optimal combination strategy we identified was often only marginally better than the PD03525901+PLX-4720 gold-standard reference. Similarly, for the multi-cell line analysis, the algorithm identified the gold-standard combination for low total dosages and was only able to identify better combinations at higher dosages. However, we consistently observed high concentrations of the MEK inhibitor PD0325901, which is known to display otherwise rare on-target toxicities, suggesting that a different regularization strategy might be desired for this drug.

One limitation of the current study is that the relative cell viability measures we have used here, such as those reported by assays such as CellTiter-Glo, are subject to several known inconsistencies [19, 24]. These issues can, in part, be addressed by more modern methods [18, 37]. Similarly, the assumption that cell growth dynamics have reached a steady-state after 72 hours may not always hold true. This may influence whether and how well biological insights presented in this study can be replicated in *in vitro* and *in vivo* experiments. However, these limitations are primarily due to limitations of data available in the large pharmacological studies [4] that were used in the parameterization of the current model, and not due to intrinsic shortcomings of the methods developed in this study. In fact, the methods developed here could easily be applied to the design of adaptive treatment strategies [51].

The model employed here assumes cell-line-specific, but static transcription. Accordingly, the model may not accurately describe adaptive resistance mechanisms that are believed to work through transcriptional feedbacks [17, 11]. Moreover, because the steady state of the model is always unimodal under conditions we have considered, there is no memory effect between subsequent treatments at the cellular level. However, the multiplicative propagation of relative viabilities along the sequence of treatments introduces a memory effect at the population level. In every treatment step, the relative proliferation values from the previous step effectively introduce a re-weighting of the relative importance of the cell lines. As we showed, this alone is enough to cause there to be benefit from time-varying sequential treatments. In practice, a further benefit from sequential treatment may be obtainable by steering a cell line or set of cell lines during the dynamics, that is, without waiting for steady state between treatments. Finding such treatment plans computationally would require a signaling pathway model that is faithful to reality not only at steady states but also during the transient paths. Constructing and calibrating such models would likely require significantly more *in vivo* and/or *in vitro* data than models that only need to be accurate in steady states.

For some cell-lines and regularizers, we observed that optimization can yield a continuum of equivalent optimal treatments, which indicates ill-conditioning of the problem. Further investigation (Appendix A.3) revealed that this behaviour is limited to low penalization strengths that do not reduce the total concentration of the optimal treatment beyond the 8 *µM* maximum. Accordingly, we concluded that this ill-conditioning did not substantially effect the results present here and that the regularization approaches, as expected, improved the conditioning of the problem.

The regularization functions we used provide an empirical way to minimize drug concentrations and adverse toxicities. In practice, concentrations at which adverse toxicities occur may be specific to drugs, tissues, and person. In the absence of large-scale toxicological and pharmacokinetic screenings, it seems difficult to design a more rational type and strength of penalization. Our regularization functions penalize total drug burden and do not consider cooperativity in adverse toxicity.

## 5 Conclusions

In this paper we proposed a framework for *in silico* combination treatment optimization. To the best of our knowledge this is the first time a large-scale pan-cancer pathway model was used to identify effective treatment strategies. Multiple treatment optimization problems were proposed which required us to balance reduction in proliferation with adverse side effects. In order to solve these problems, we combined the CMA-ES algorithm with a significantly more scalable sampling scheme, based on a Hamiltonian Monte-Carlo method. We evaluated the approach in an extensive simulation study with cancer cell lines originating from multiple tissues. We studied the treatment of individual cell lines and heterogeneous populations of cell lines. We also studied the generation of sequential time-varying and time-invariant treatment plans. The combination treatments identified by our algorithm achieved significantly better predicted proliferation scores at lower drug concentrations compared to the conventional therapy approaches. This serves as an early proof of concept of how *in silico* simulations can be used to identify potentially novel combination therapies. Future research is required to evaluate the performance of the discovered treatments in laboratory studies.

## 6 Broader Impact

The field of systems biology strives to understand and model complex biological systems through means of computational and mathematical analysis. Recent years have led to increasingly sophisticated models such as the pan-cancer pathway model used in this paper, which are the result of rich domain knowledge and large amounts of experimental data. In this context our work provides a methodology to extract the information encoded into these models to gain actionable insights in the form of potentially novel combination therapies. One can imagine a not-too-far-away future in which techniques like the ones proposed in this paper are used to perform large-scale *in silico* experiments to identify a small set of promising treatment candidates. The treatments that have proven themselves in simulations can then be used as a starting point for laboratory studies, which could lead to more efficient use of limited resources and accelerated discovery of effective therapies.

## Acknowledgements

TS, RS, and GF are supported in part by TS’s ARO awards W911NF-17-1-0082 and W911NF2010081 and TS’s NSF grants IIS-1718457, IIS-1617590, IIS-1901403, and CCF-1733556. TS is Founder, President, and CEO of Sandholm Enterprises, Ltd., Strategic Machine, Inc., Strategy Robot, Inc., and Optimized Markets, Inc.; these affiliations did not affect the conclusions of this paper. GF is also supported in part by a Facebook graduate student fellowship. AS and JF are supported in part by support from NIH grant P41-GM103712 for the National Center for Multiscale Modeling of Biological Systems (MMBioS). FF is supported by HFSP grant LT000259/2019-L1 and NIH grant U54-CA225088. The conclusions of the paper are those of the authors and may not represent views of their funding agencies or organizations.

## A Supplementary Information

### A.1 Exponential Growth Model

For each cell line the relative proliferation value achieved by a sequential treatment plan is computed by taking the product of the predicted relative proliferation values at the individual treatment steps. A simple, biologically plausible model that satisfies this assumption is an exponential growth model with different, drug-dependent growth rates in each treatment step. Such a model is of the form

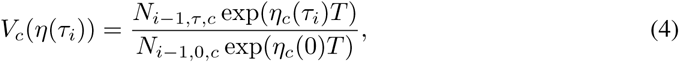

where *N* − _*i, τ,c*_ is the final cell count of cell line *c* from the previous step in the treated condition, *N*_*i* −1.0,*c*_ is the final cell count of cell line *c* from the previous step *i* − 1 in the untreated condition, *η*_*c*_(*τ*_*i*_) is the treatment-dependent growth rate of cell line *c* during the current step *i*, exp(*η*_*c*_ (0)) is the untreated growth rate of cell line *c*, and *T* is the treatment duration (which we assume to be 72 hours, the time used to generate experimental data the pathway model was calibrated on in prior work [13]). Under the assumption of such an exponential growth model, the following equations hold at every treatment step:

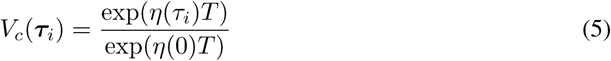

and, by induction,

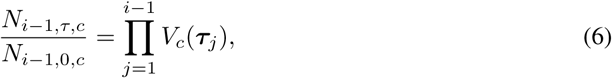

assuming that,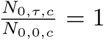 that is, both treated and untreated cell populations start at the same cell counts. This was true for the experimental data used for training the model in prior work [4] and matches the assumptions of our simulation study.

### A.2 Optimization for Average Proliferation

While this paper mainly focused on minimizing the maximum proliferation value in populations of cancer cell lines, we will now briefly discuss experimental results where the objective was to minimize the average proliferation value. This alternative optimization problem was discussed in Subsection 2.2.2.

First we investigated what average proliferation rate the multi-cell combination treatments that were optimized for the *maximum* criterion achieve for each of the four individual tissues under L1 regularization with penalty value *λ* = 10^−7^. We found that the treatments optimized for low maximum proliferation achieve an average relative proliferation rate of approximately 0.1036 across melanoma, 0.0814 across colorectal, 0.1019 across pancreatic, and 0.2010 across breast cancer cell lines. We compared these scores to those attained by the multi-cell combination treatments which were specifically designed to minimize the *average* proliferation rate. We found that the treatments optimized for low average proliferation rate achieve average proliferation values of approximately 0.0816 across melanoma, 0.0712 across colorectal, 0.0804 across pancreatic, and 0.1589 across breast cancers. Therefore, the multi-cell treatments considered in this paper not only minimize the maximum proliferation rate of cells originating from each tissue type, but they also attain average proliferation rates that are experimentally within 20% of what is attained by the treatments which were specifically designed for low average.

### A.3 Variance in Optimization Process

During our single- and multi-cell experiments (Section 3) we observed some variance in the optimized combination treatments when using low penalty values. This indicates the existence of multiple local optima. To get a better insight into this behavior we performed an additional single-cell experiment with K029AX as well as a multi-cell experiment with colorectal cell lines. For both settings we ran an additional 20 runs with warm starts. Each run started by optimizing a treatment for the lowest penalty value (10^−7^) and then increased the penalty exponent at 0.25 steps, where at each step we initialized the algorithm with the optimal drug-combination from the previous step.

We grouped the discovered drug-combinations found during the 20 runs by penalty value and performed separate Principal Component Analysis (PCA) for each group to investigate the treatment distribution. The first two principal components are visualized in Figure 5 and 6 which in both experiments explained more than 90% of the existing variance. Under high to medium penalties L1 and L2 regularization led to unique optimal treatments. For lower penalty values there is some variance. Logarithmic penalization suffers from high variance even when using large penalties indicating many local optima. This might explain some of the instabilities we observed in the previous experiments which used logarithmic regularization. For low penalty values the distribution of the returned combinations is similar for all types of regularization. Overall the variance in the multi-cell experiment is larger than in the single-cell one.

**Figure 5:**
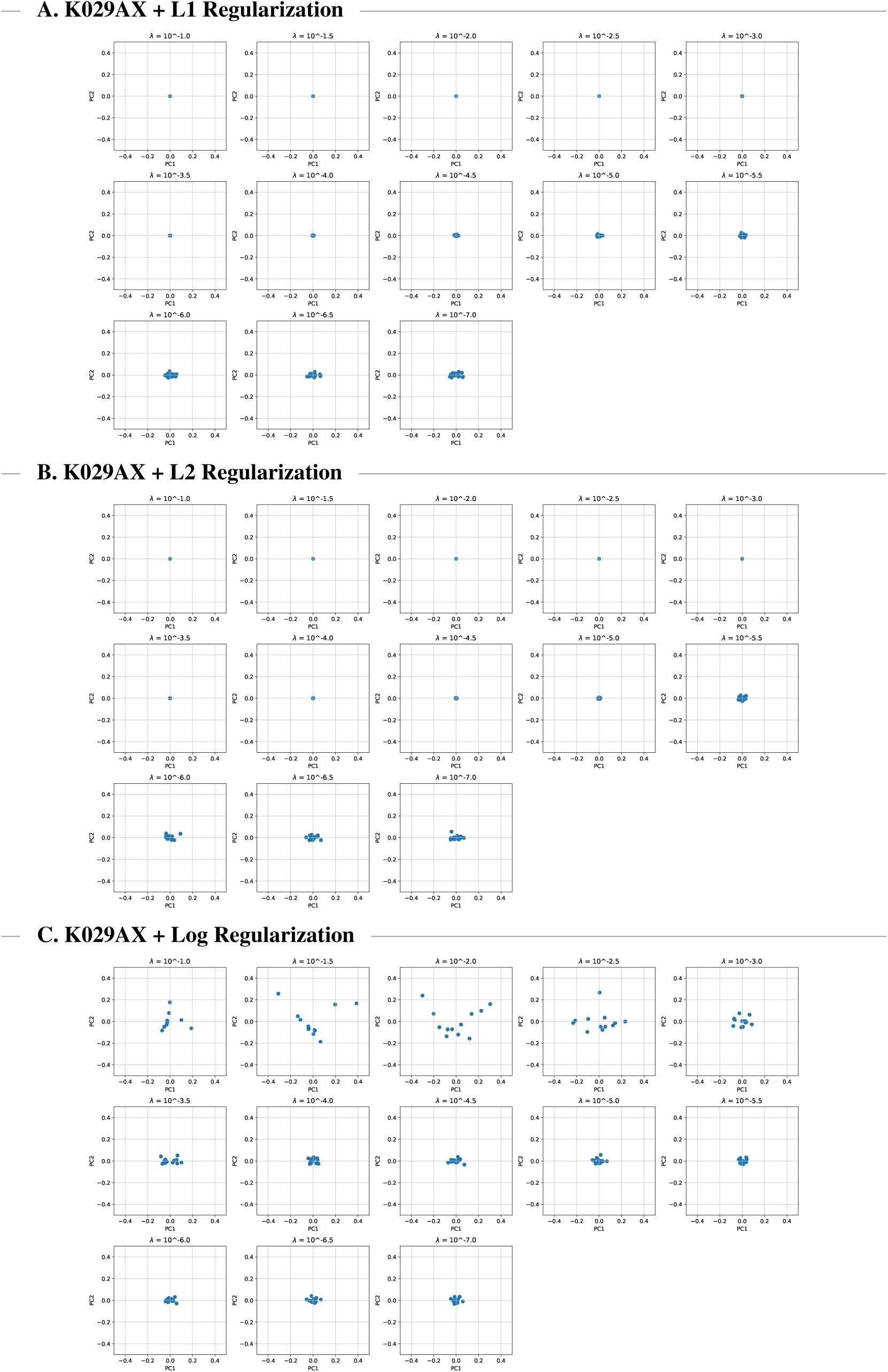
Visualization of the first two principal components of 20 single-cell combination treatments for K029AX under three different types of regularization using warm starts.

**Figure 6:**
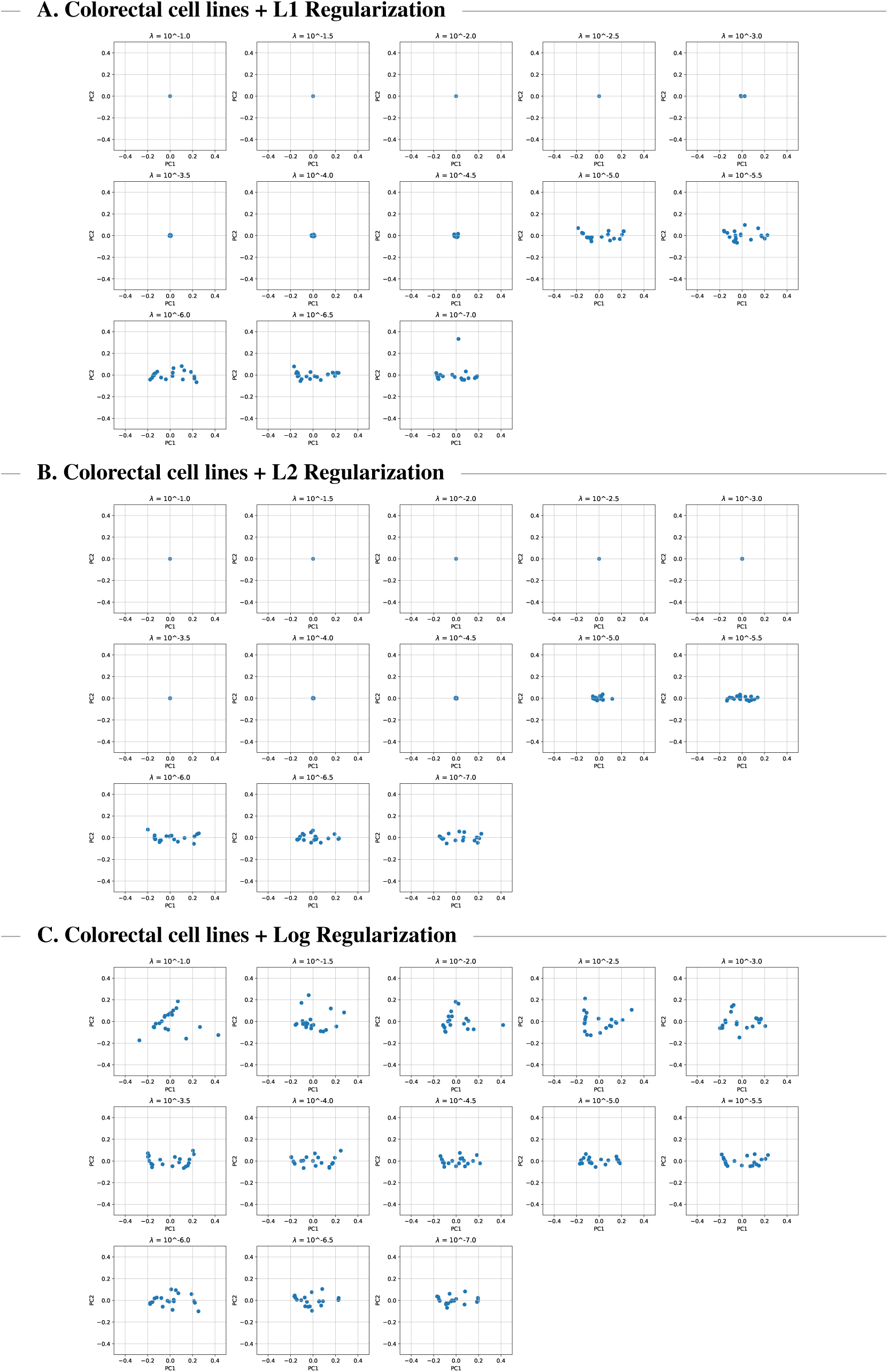
Visualization of the first two principal components of 20 multi-cell combination treatments for colorectal cells under three different types of regularization using warm starts.

### A.4 Further Single-Step Single Cell Experiments

Figure 7 and 8 show the results of the single-step single-cell optimization process for A2058 and MDAMB43S cells respectively. For A2058 we observed that for all three types of regularization the optimized combination treatments achieve significantly lower relative proliferation values at lower concentrations than the single and two-drug baselines. For MDAMB43S the discovered combination treatments only slightly improved upon the PD0325901/PLX-4720 two-drug baseline. In both cases the type of regularization impacts the composition of the returned combination treatments. When using logarithmic regularization we observed large variance in returned treatments and low objective values did not always indicate effective treatments. This behaviour is further investigated in Appendix A.3.

**Figure 7:**
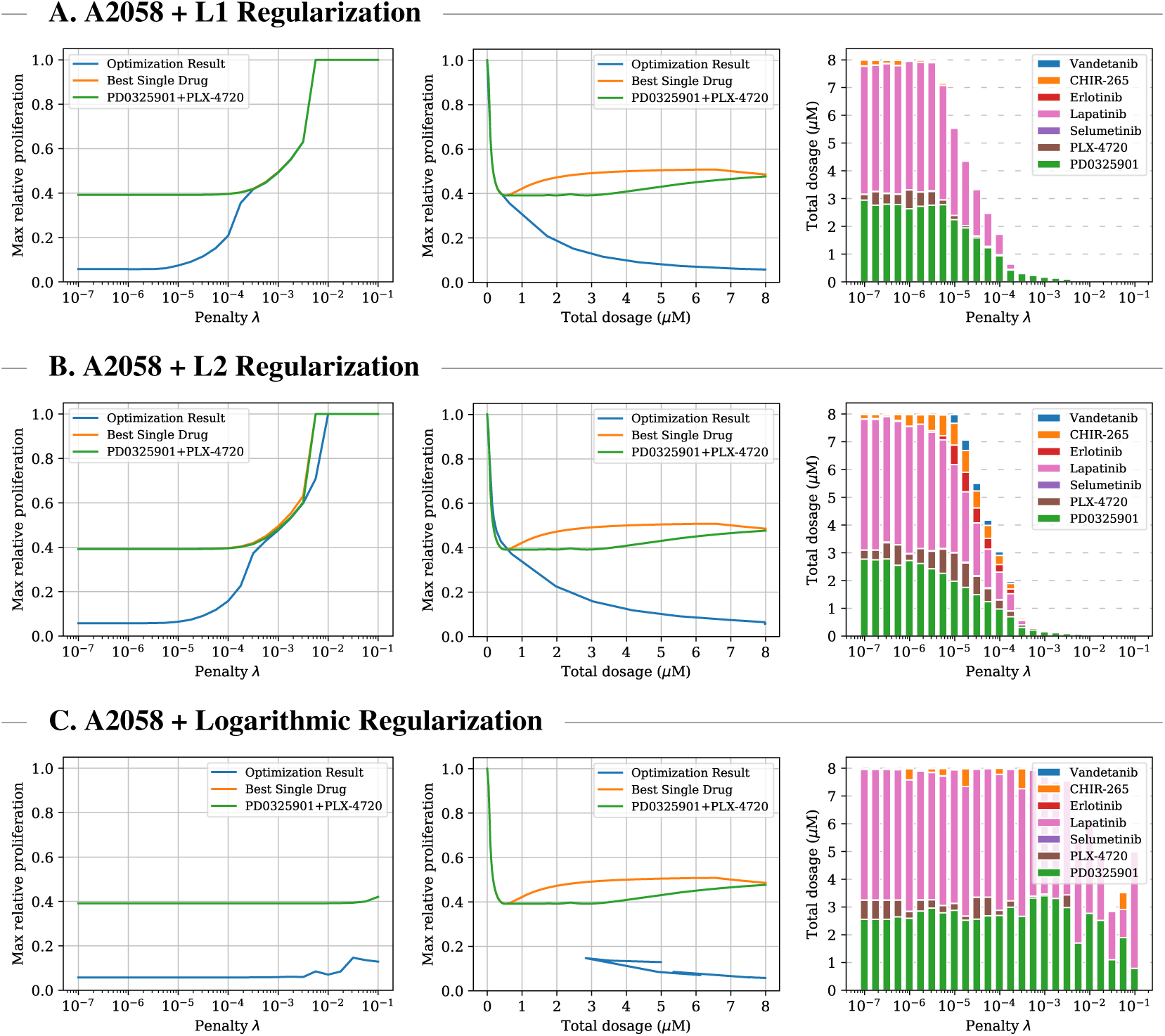
Comparison between optimized single-cell multi-drug treatment, optimal single-drug treatment, and optimal PD0325901/PLX-4720 combination treatment for A2058—a melanoma cell line with BRAF V600E mutation—for three different types of regularization. Left plots: optimal treatment as identified by the objective function for different penalty parameters. The middle plots: relationship between administered total dosage and achieved proliferation value regardless of penalty and objective value. Right plots: composition of the multi-drug treatments.

**Figure 8:**
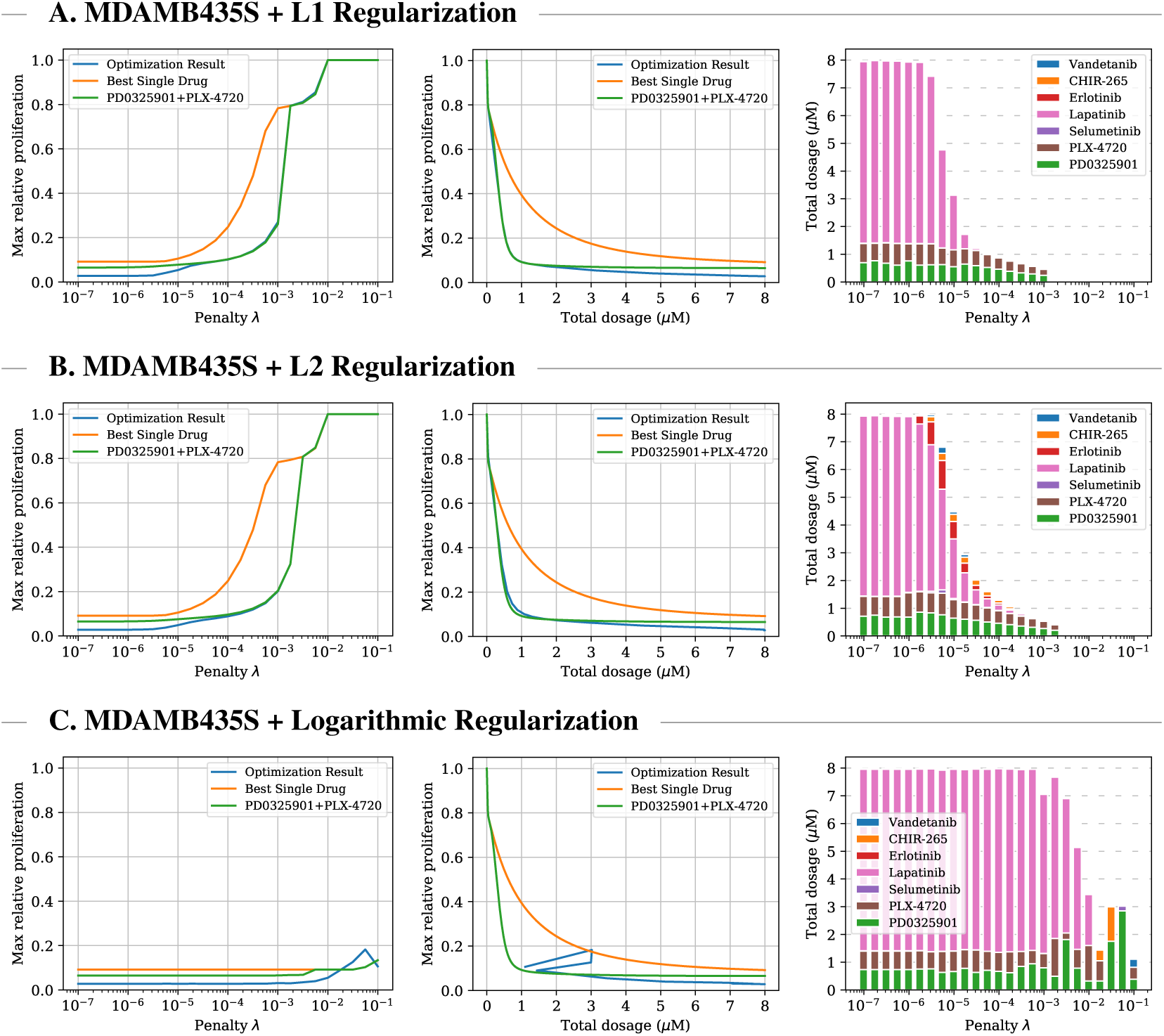
Comparison between optimized single-cell multi-drug treatment, optimal single-drug treatment, and optimal PD0325901/PLX-4720 combination treatment for MDAMB435S—a melanoma cell line with BRAF V600E mutation—for three different types of regularization. Left plots: optimal treatment as identified by the objective function for different penalty parameters. The middle plots: relationship between administered total dosage and achieved proliferation value regardless of penalty and objective value. Right plots: composition of the multi-drug treatments.

### A.5 Further Single-Step Population Cell Experiments

We show results of the single-step multi-cell optimization process for melanoma (Figure 9), colorectal (Figure 10), pancreatic (Figure 11) and breast (Figure 12) cancer cell lines. For all four tissues and regularizers, the discovered combination treatments achieve significantly lower maximum relative proliferation values than the single-drug and PD0325901/PLX-4720 combination baselines at medium and high dosages. Especially for pancreatic cell lines, the optimized treatments reduce the cancer cell viability by a factor of more than two. For breast cancers, the optimization process leads to drug combinations that achieve notable treatment effects even at low dosage. The type of used regularization effects the composition of the combinations. When using logarithmic regularization we observed large variance in returned treatments and low objective values did not always indicate effective treatments. This behaviour is further investigated in Appendix A.3.

**Figure 9:**
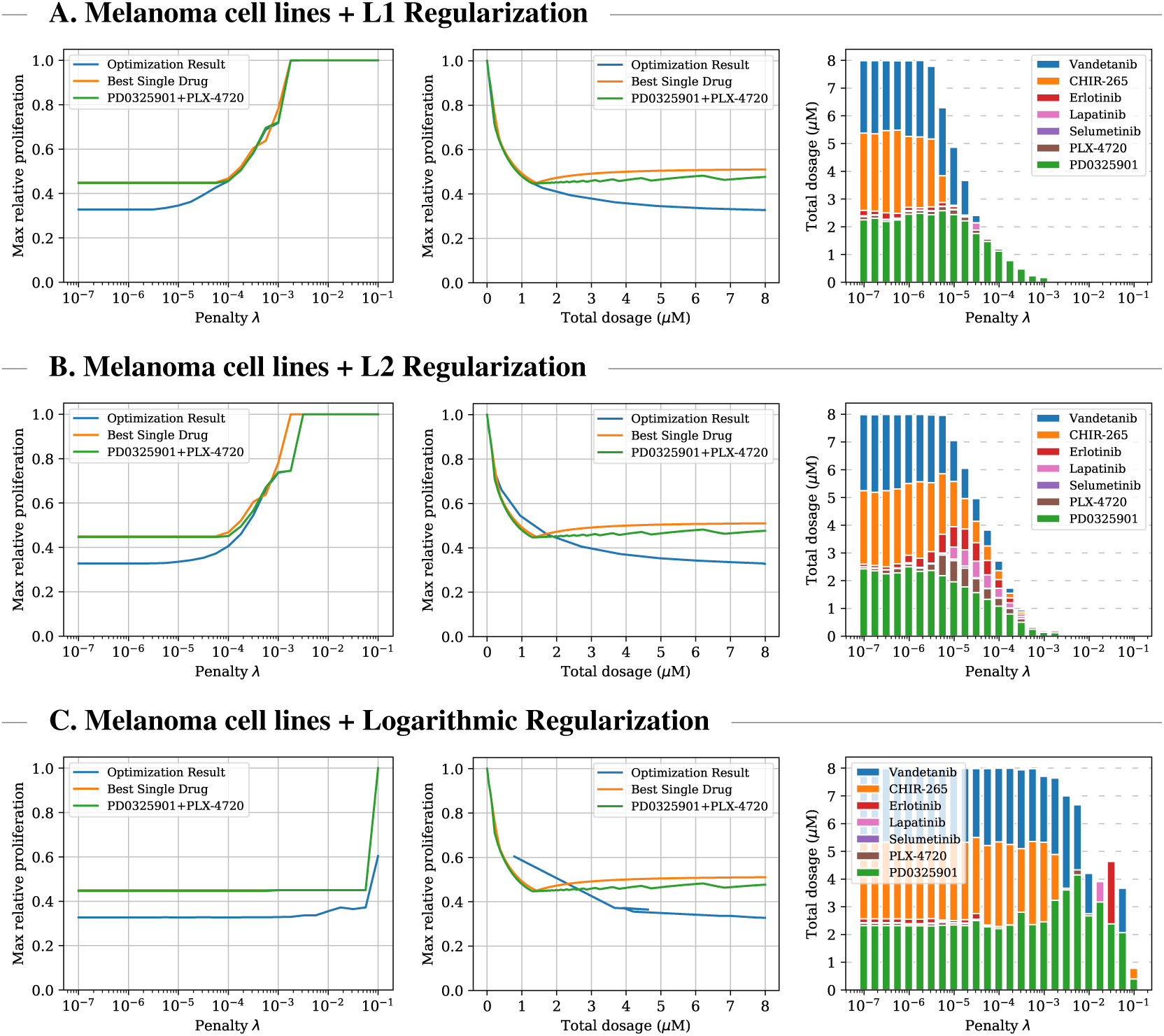
Comparison between optimized multi-cell multi-drug treatment, optimal single-drug treatment, and optimal PD0325901/PLX-4720 combination treatment for melanoma cell lines for three different types of regularization. Left plot: optimal treatment as identified by the objective function for different penalty parameters. Middle plot: relationship between administered total dosage and achieved proliferation value regardless of objective values. Right plot: composition of the multi-drug treatments.

**Figure 10:**
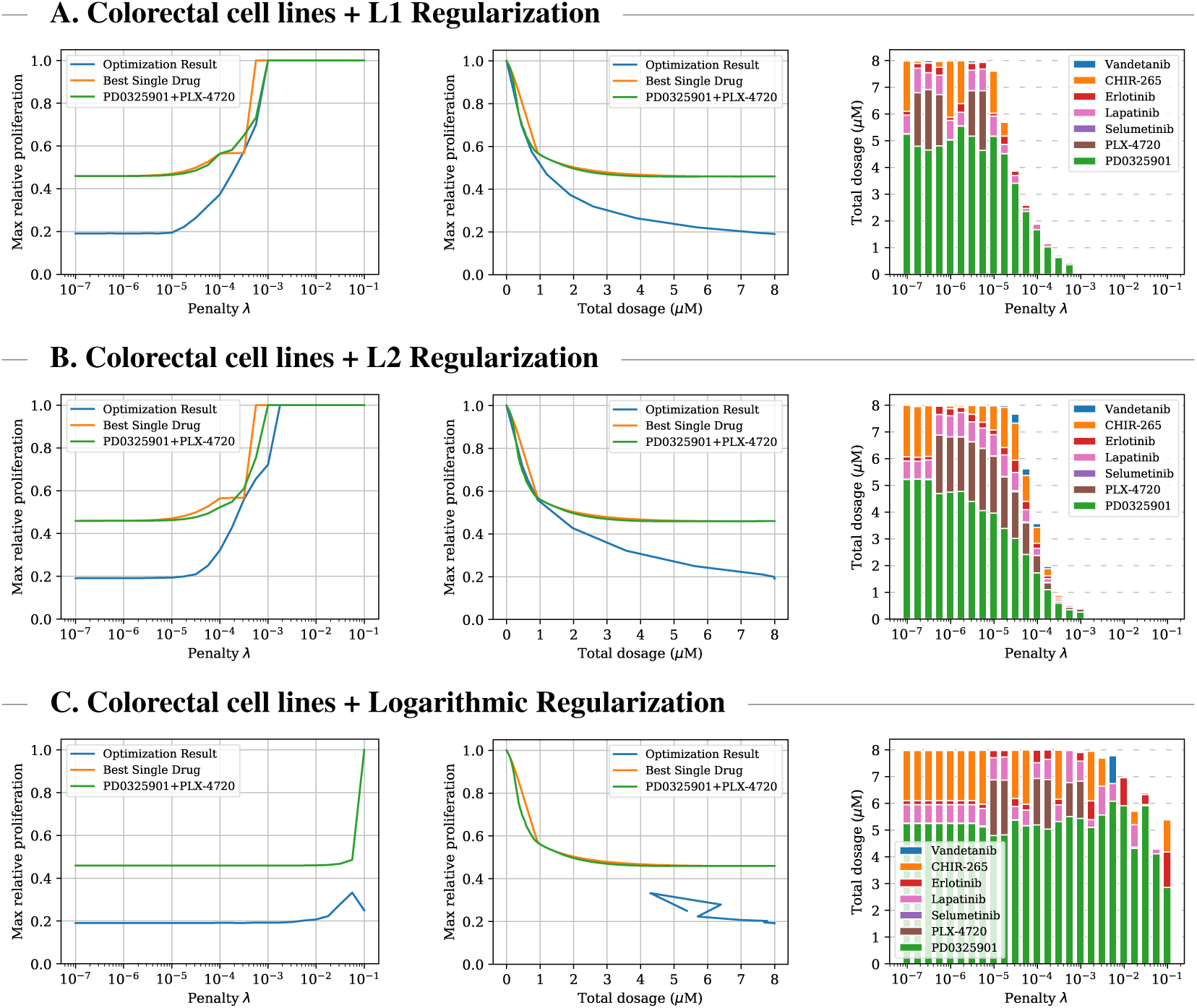
Comparison between optimized multi-cell multi-drug treatment, optimal single-drug treatment, and optimal PD0325901/PLX-4720 combination treatment for colorectal cell lines for three different types of regularization. Left plot: optimal treatment as identified by the objective function for different penalty parameters. Middle plot: relationship between administered total dosage and achieved proliferation value regardless of objective values. Right plot: composition of the multi-drug treatments.

**Figure 11:**
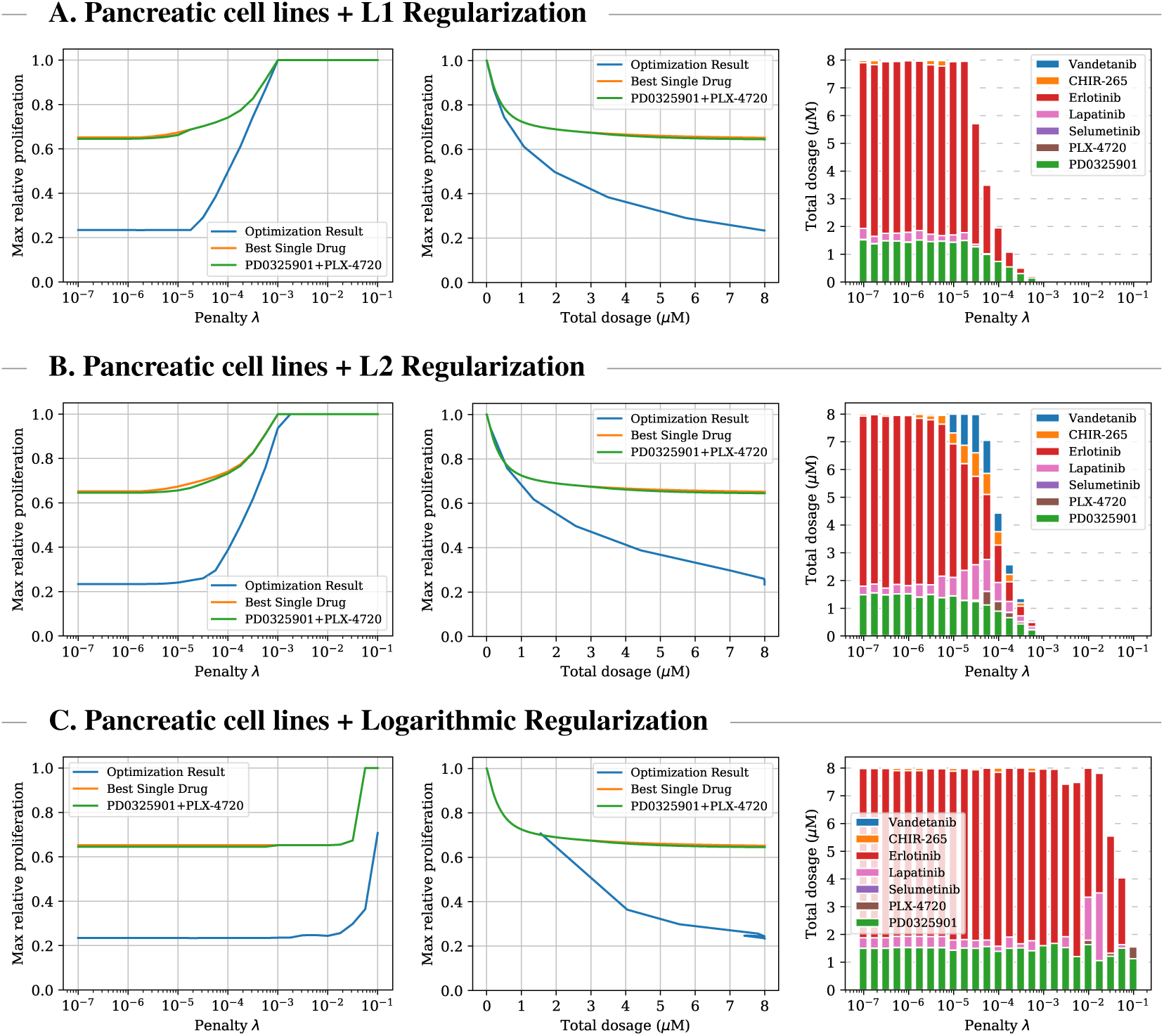
Comparison between optimized multi-cell multi-drug treatment, optimal single-drug treatment, and optimal PD0325901/PLX-4720 combination treatment for pancreatic cell lines for three different types of regularization. Left plot: optimal treatment as identified by the objective function for different penalty parameters. Middle plot: relationship between administered total dosage and achieved proliferation value regardless of objective values. Right plot: composition of the multi-drug treatments.

**Figure 12:**
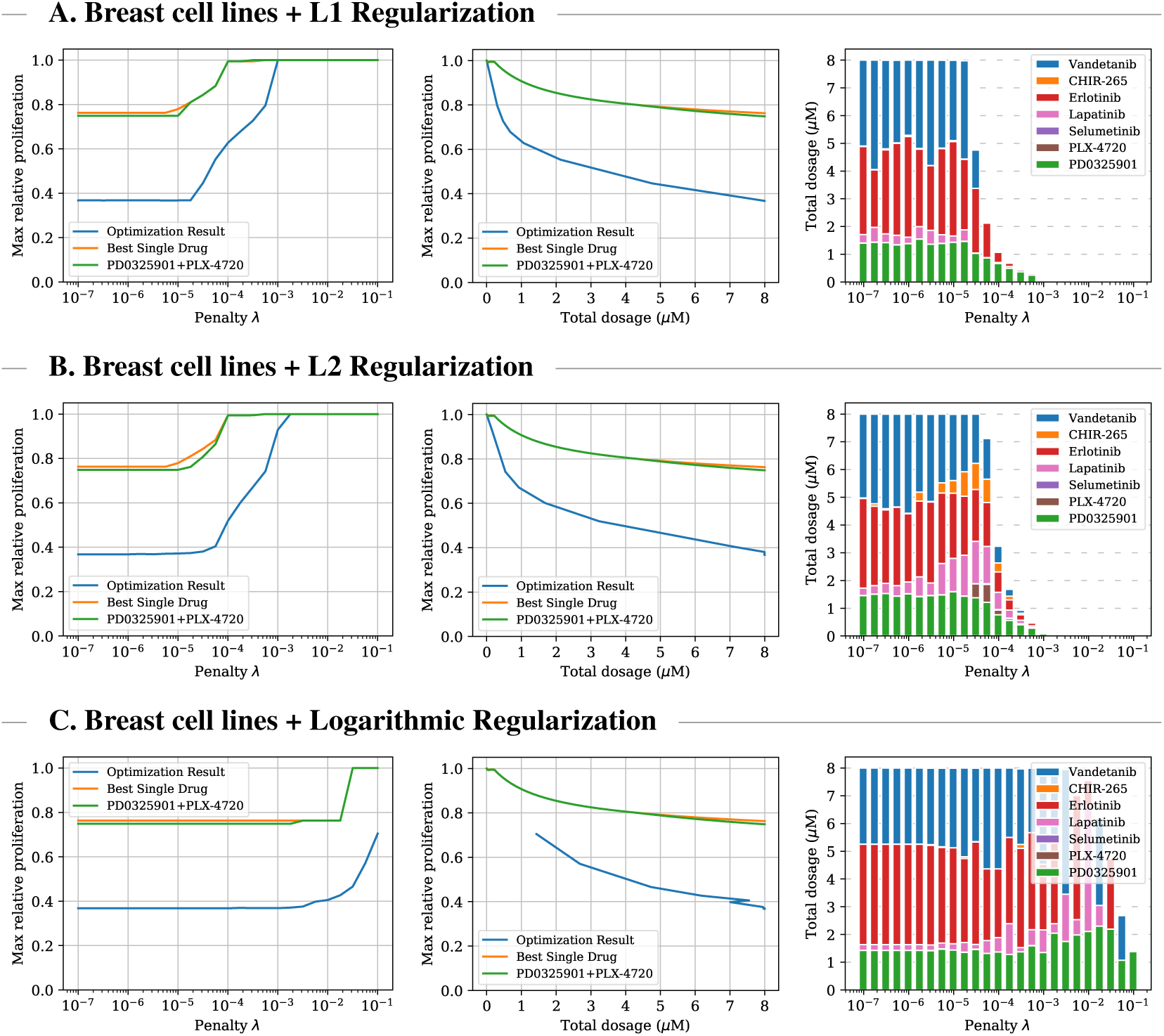
Comparison between optimized multi-cell multi-drug treatment, optimal single-drug treatment, and optimal PD0325901/PLX-4720 combination treatment for breast cancer cell lines for three different types of regularization. Left plot: optimal treatment as identified by the objective function for different penalty parameters. Middle plot: relationship between administered total dosage and achieved proliferation value regardless of objective values. Right plot: composition of the multi-drug treatments.

### A.6 Numerical Results of Sequential Treatment Experiment

We provide numerical results for the previous two-step treatment optimization experiment visualized in Figure 4. Figure 13 compares the relative proliferation values achieved by the drug cocktails administered by the optimized two-step treatment plan and the optimized two-step time-invariant treatment plan for colorectal cell lines at the same total drug dosage (550 nM). The 2-step plan uses a high-dose treatment followed by a low-dose treatment. This achieves maximum proliferation 0.6048, which is more effective than the time-invariant treatment plan which only achieves 0.6978. The table shows the relative proliferation values for each individual colorectal cell lines after the first and second treatment step. The highest proliferation values after each treatment step are marked bold.

**Figure 13:**
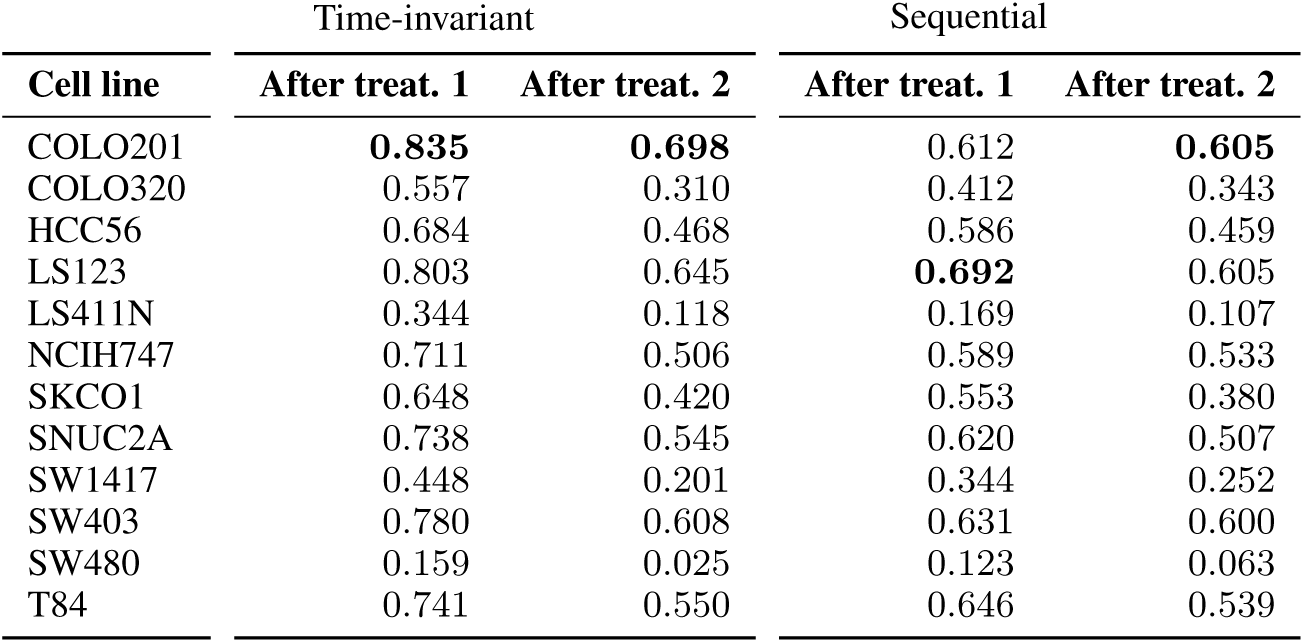
Comparison of the relative proliferation values achieved by the drug cocktails administered by the optimized two-step treatment plan and the optimized two-step time-invariant treatment plan for colorectal cell lines at the same total drug dosage (550 nM) after first and second step. The employed treatments are described by Figure 4.

## References

[1] Ahmet Acar, Daniel Nichol, Javier Fernandez-Mateos, George D Cresswell, Iros Barozzi, Sung Pil Hong, Nicholas Trahearn, Inmaculada Spiteri, Mark Stubbs, Rosemary Burke, et al. Exploiting evolutionary steering to induce collateral drug sensitivity in cancer. Nature Communications, 11(1):1–14, 2020.

[2] Bissan Al-Lazikani, Udai Banerji, and Paul Workman. Combinatorial drug therapy for cancer in the post-genomic era. Nature biotechnology, 30(7):679, 2012.

[3] K Bahrami and M Kim. Optimal control of multiplicative control systems arising from cancer therapy. IEEE Transactions on Automatic Control, 20(4):537–542, 1975.

[4] Jordi Barretina, Giordano Caponigro, Nicolas Stransky, Kavitha Venkatesan, Adam A Margolin, Sungjoon Kim, Christopher J Wilson, Joseph Lehár, Gregory V Kryukov, Dmitriy Sonkin, et al. The cancer cell line encyclopedia enables predictive modelling of anticancer drug sensitivity. Nature, 483(7391):603–607, 2012.

[5] Rafal Biedrzycki. Handling bound constraints in CMA-ES: An experimental study. Swarm and Evolutionary Computation, 52:100627, 2020.

[6] Mehdi Bouhaddou, Anne Marie Barrette, Alan D Stern, Rick J Koch, Matthew S DiStefano, Eric A Riesel, Luis C Santos, Annie L Tan, Alex E Mertz, and Marc R Birtwistle. A mechanistic pan-cancer pathway model informed by multi-omics data interprets stochastic cell fate responses to drugs and mitogens. PLoS computational biology, 14(3):e1005985, 2018.

[7] Cécile Carrere. Optimization of an in vitro chemotherapy to avoid resistant tumours. Journal of Theoretical Biology, 413:24–33, 2017.

[8] Federica Eduati, Victoria Doldàn-Martelli, Bertram Klinger, Thomas Cokelaer, Anja Sieber, Fiona Kogera, Mathurin Dorel, Mathew J Garnett, Nils Blüthgen, and Julio Saez-Rodriguez. Drug resistance mechanisms in colorectal cancer dissected with cell type–specific dynamic logic models. Cancer research, 77(12):3364–3375, 2017.

[9] Pedro M Enriquez-Navas, Yoonseok Kam, Tuhin Das, Sabrina Hassan, Ariosto Silva, Parastou Foroutan, Epifanio Ruiz, Gary Martinez, Susan Minton, Robert J Gillies, et al. Exploiting evolutionary principles to prolong tumor control in preclinical models of breast cancer. Science translational medicine, 8(327):327ra24–327ra24, 2016.

[10] Warren J Ewens. Mathematical population genetics 1: Theoretical Introduction, volume 27. Springer Science & Business Media, 2012.

[11] Mohammad Fallahi-Sichani, Verena Becker, Benjamin Izar, Gregory J Baker, Jia-Ren Lin, Sarah A Boswell, Parin Shah, Asaf Rotem, Levi A Garraway, and Peter K Sorger. Adaptive resistance of melanoma cells to raf inhibition via reversible induction of a slowly dividing de-differentiated state. Molecular systems biology, 13(1), 2017.

[12] Fabian Fröhlich, Barbara Kaltenbacher, Fabian J Theis, and Jan Hasenauer. Scalable parameter estimation for genome-scale biochemical reaction networks. PLoS computational biology, 13(1), 2017.

[13] Fabian Fröhlich, Thomas Kessler, Daniel Weindl, Alexey Shadrin, Leonard Schmiester, Hendrik Hache, Artur Muradyan, Moritz Schütte, Ji-Hyun Lim, Matthias Heinig, et al. Efficient parameter estimation enables the prediction of drug response using a mechanistic pan-cancer pathway model. Cell Systems, 7(6):567–579, 2018.

[14] Jill A Gallaher, Pedro M Enriquez-Navas, Kimberly A Luddy, Robert A Gatenby, and Alexander RA Anderson. Spatial heterogeneity and evolutionary dynamics modulate time to recurrence in continuous and adaptive cancer therapies. Cancer research, 78(8):2127–2139, 2018.

[15] Robert A Gatenby, Ariosto S Silva, Robert J Gillies, and B Roy Frieden. Adaptive therapy. Cancer research, 69(11):4894–4903, 2009.

[16] Robert A Gatenby and Thomas L Vincent. Application of quantitative models from population biology and evolutionary game theory to tumor therapeutic strategies. Molecular cancer therapeutics, 2(9):919–927, 2003.

[17] Luca Gerosa, Christopher Chidley, Fabian Froehlich, Gabriela Sanchez, Sang Kyun Lim, Jeremy Muhlich, Jia-Yun Chen, Gregory J Baker, Denis Schapiro, Tujin Shi, et al. Sporadic erk pulses drive non-genetic resistance in drug-adapted brafv600e melanoma cells. bioRxiv, page 762294, 2019.

[18] Marc Hafner, Mario Niepel, Mirra Chung, and Peter K Sorger. Growth rate inhibition metrics correct for confounders in measuring sensitivity to cancer drugs. Nature methods, 13(6):521, 2016.

[19] Benjamin Haibe-Kains, Nehme El-Hachem, Nicolai Juul Birkbak, Andrew C Jin, Andrew H Beck, Hugo JWL Aerts, and John Quackenbush. Inconsistency in large pharmacogenomic studies. Nature, 504(7480):389–393, 2013.

[20] Rita Hannah, Michael Beck, Richard Moravec, and T Riss. Celltiter-gloTM luminescent cell viability assay: a sensitive and rapid method for determining cell viability. Promega Cell Notes, 2:11–13, 2001.

[21] Nikolaus Hansen. The CMA evolution strategy: A tutorial. arXiv preprint arXiv:1604.00772, 2016.

[22] Nikolaus Hansen and Andreas Ostermeier. Completely derandomized self-adaptation in evolution strategies. Evol. Comput., 9(2):159–195, June 2001.

[23] A. Hastings. Population Biology: Concepts and Models. Ecology. Mathematical biology). Springer New York, 1996.

[24] Peter M Haverty, Eva Lin, Jenille Tan, Yihong Yu, Billy Lam, Steve Lianoglou, Richard M Neve, Scott Martin, Jeff Settleman, Robert L Yauch, et al. Reproducible pharmacogenomic profiling of cancer cell line panels. Nature, 533(7603):333–337, 2016.

[25] W F Hawse, R P Sheehan, Natasa Miskov-Zivanov, A V Menk, L P Kane, James R Faeder, and Penelope A Morel. Cutting edge: Differential regulation of PTEN by TCR, Akt, and FoxO1 controls CD4+ T cell fate decisions. J Immunol, 194:4615–4619, 2015.

[26] Alan C Hindmarsh, Peter N Brown, Keith E Grant, Steven L Lee, Radu Serban, Dan E Shumaker, and Carol S Woodward. Sundials: Suite of nonlinear and differential/algebraic equation solvers. ACM Transactions on Mathematical Software (TOMS), 31(3):363–396, 2005.

[27] Josef Hofbauer, Karl Sigmund, et al. Evolutionary games and population dynamics. Cambridge university press, 1998.

[28] Mykel J Kochenderfer and Tim A Wheeler. Algorithms for optimization. MIT Press, 2019.

[29] Walter Kolch and Dirk Fey. Personalized computational models as biomarkers. Journal of personalized medicine, 7(3):9, 2017.

[30] Christian Kroer, Gabriele Farina, and Tuomas Sandholm. Robust Stackelberg equilibria in extensive-form games and extension to limited lookahead. In AAAI Conference on Artificial Intelligence (AAAI), 2018.

[31] Christian Kroer and Tuomas Sandholm. Limited lookahead in imperfect-information games. In Proceedings of the International Joint Conference on Artificial Intelligence (IJCAI), 2015.

[32] Christian Kroer and Tuomas Sandholm. Sequential planning for steering immune system adaptation. In Proceedings of the International Joint Conference on Artificial Intelligence (IJCAI), IJCAI’16, page 3177–3184. AAAI Press, 2016.

[33] James Larkin, Paolo A Ascierto, Brigitte Dréno, Victoria Atkinson, Gabriella Liszkay, Michele Maio, Mario Mandalà, Lev Demidov, Daniil Stroyakovskiy, Luc Thomas, et al. Combined vemurafenib and cobimetinib in braf-mutated melanoma. New England Journal of Medicine, 371(20):1867–1876, 2014.

[34] Urszula Ledzewicz and Heinz Schättler. Drug resistance in cancer chemotherapy as an optimal control problem. Discrete & Continuous Dynamical Systems-B, 6(1):129, 2006.

[35] João M Lemos, Daniela V Caiado, Rui Coelho, and Susana Vinga. Optimal and receding horizon control of tumor growth in myeloma bone disease. Biomedical Signal Processing and Control, 24:128–134, 2016.

[36] Jieru Meng, Bingbing Dai, Bingliang Fang, B Nebiyou Bekele, William G Bornmann, Duoli Sun, Zhenghong Peng, Roy S Herbst, Vassiliki Papadimitrakopoulou, John D Minna, et al. Combination treatment with mek and akt inhibitors is more effective than each drug alone in human non-small cell lung cancer in vitro and in vivo. PloS one, 5(11), 2010.

[37] Mario Niepel, Marc Hafner, Caitlin E Mills, Kartik Subramanian, Elizabeth H Williams, Mirra Chung, Benjamin Gaudio, Anne Marie Barrette, Alan D Stern, Bin Hu, et al. A multi-center study on the reproducibility of drug-response assays in mammalian cell lines. Cell systems, 9(1):35–48, 2019.

[38] Elin Nyman, Richard R Stein, Xiaohong Jing, Weiqing Wang, Benjamin Marks, Ioannis K Zervantonakis, Anil Korkut, Nicholas P Gauthier, and Chris Sander. Perturbation biology links temporal protein changes to drug responses in a melanoma cell line. BioRxiv, page 568758, 2019.

[39] Paul A Orlando, Robert A Gatenby, and Joel S Brown. Cancer treatment as a game: Integrating evolutionary game theory into the optimal control of chemotherapy. Physical biology, 9(6):065007, 2012.

[40] Ari Pakman and Liam Paninski. Exact Hamiltonian Monte Carlo for truncated multivariate Gaussians. Journal of Computational and Graphical Statistics, 23(2):518–542, 2014.

[41] Oleksii S Rukhlenko, Fahimeh Khorsand, Aleksandar Krstic, Jan Rozanc, Leonidas G Alexopoulos, Nora Rauch, Keesha E Erickson, William S Hlavacek, Richard G Posner, Silvia Gómez-Coca, et al. Dissecting raf inhibitor resistance by structure-based modeling reveals ways to overcome oncogenic ras signaling. Cell systems, 7(2):161–179, 2018.

[42] Julio Saez-Rodriguez and Nils Blüthgen. Personalized signaling models for personalized treatments. Molecular Systems Biology, 16(1), 2020.

[43] Tuomas Sandholm. Steering evolution strategically: Computational game theory and opponent exploitation for treatment planning, drug design, and synthetic biology. In AAAI Conference on Artificial Intelligence (AAAI), AAAI’15, page 4057–4061. AAAI Press, 2015.

[44] Tuomas W Sandholm. Medical treatment planning via sequential games, 2012. Provisional patent application. Converted to full Patent Application 13/955,966.

[45] Afroza Shirin, Isaac S Klickstein, Song Feng, Yen Ting Lin, William S Hlavacek, and Francesco Sorrentino. Prediction of optimal drug schedules for controlling autophagy. Scientific reports, 9(1):1–15, 2019.

[46] John Maynard Smith. Evolution and the Theory of Games. Cambridge university press, 1982.

[47] Kateřina Staňková, Joel S Brown, William S Dalton, and Robert A Gatenby. Optimizing cancer treatment using game theory. JAMA oncology, 5(1):96–103, 2019.

[48] George W Swan and Thomas L Vincent. Optimal control analysis in the chemotherapy of igg multiple myeloma. Bulletin of mathematical biology, 39(3):317–337, 1977.

[49] Andrzej Swierniak, Marek Kimmel, and Jaroslaw Smieja. Mathematical modeling as a tool for planning anticancer therapy. European journal of pharmacology, 625(1-3):108–121, 2009.

[50] Jeffrey West, Mark Robertson-Tessi, Kimberly Luddy, Derek S Park, Drew FK Williamson, Cathal Harmon, Hung T Khong, Joel Brown, and Alexander RA Anderson. The immune checkpoint kick start: Optimization of neoadjuvant combination therapy using game theory. JCO clinical cancer informatics, 3:1–12, 2019.

[51] Jeffrey West, Li You, Jingsong Zhang, Robert A Gatenby, Joel S Brown, Paul K Newton, and Alexander RA Anderson. Towards multi-drug adaptive therapy. Cancer Research, 2020.

[52] Jeffrey B West, Mina N Dinh, Joel S Brown, Jingsong Zhang, Alexander R Anderson, and Robert A Gatenby. Multidrug cancer therapy in metastatic castrate-resistant prostate cancer: an evolution-based strategy. Clinical Cancer Research, 25(14):4413–4421, 2019.

[53] Bo Yuan, Ciyue Shen, Augustin Luna, Anil Korkut, Debora S Marks, John Ingraham, and Chris Sander. Interpretable machine learning for perturbation biology. bioRxiv, page 746842, 2019.

[54] Jingsong Zhang, Jessica J Cunningham, Joel S Brown, and Robert A Gatenby. Integrating evolutionary dynamics into treatment of metastatic castrate-resistant prostate cancer. Nature communications, 8(1):1–9, 2017.

